# A “molecular guillotine” reveals an interphase function of Kinesin-5

**DOI:** 10.1101/184416

**Authors:** Zhiyi Lv, Jan Rosenbaum, Timo Aspelmeier, Jörg Großhans

## Abstract

Motor proteins are important for transport and force generation in a variety of cellular processes and morphogenesis. Here we design a general strategy for conditional motor mutants by inserting a protease cleavage site at the “neck” between the head domain and the stalk of the motor protein, making the protein susceptible to proteolytic cleavage at the neck by the corresponding protease. To demonstrate the feasibility of this approach, we inserted the cleavage site of TEV protease into the neck of the tetrameric motor Kinesin-5. Application of TEV protease led to a specific depletion and functional loss of Kinesin-5 in Drosophila embryos. By this, we revealed that Kinesin-5 stabilized the microtubule network during interphase in syncytial embryos. The “molecular guillotine” can potentially be applied to many motor proteins due to the conserved structures of kinesin, dynein and myosin with accessible necks.

**Author summary:** We design a general strategy for conditional motor mutants by inserting a protease cleavage site between head and stalk domain of the motor protein, making it susceptible to specific proteolytic cleavage. We demonstrate the feasibility of the approach with the motor Kinesin-5 and the protease TEV in Drosophila embryos. This approach can potentially be applied to motor proteins kinesin, dynein and myosin due to the conserved structures.

## Introduction

Cytoskeletal motor proteins, including myosins, dyneins and kinesins, convert the chemical energy of ATP hydrolysis into mechanical work. Motor proteins are wildly involved in multiple fundamental cellular processes such as intracellular transport, cell division, cell shape change and migration [1]. The structure of motor proteins is conserved. They contain a motor domain, referred to as “head”, which catalyzes ATP and binds microtubules or F-actin. The catalytic cycle links ATP hydrolysis to a conformational change of the protein that translates into unidirectional movement of the motor protein on the filament. A second part of the protein, the stalk, links the head to the cargo binding site, contains coil-coiled structures for oligomerization or associates with other subunits. Head and stalk are parts of the same polypeptide, which is functionally relevant as a tight link of head and stalk is essential for transmission of mechanical force [2].

Genetic analysis of the physiological function of motor proteins is hampered, since many motor proteins fulfill an essential function for the cell or organism. For example, Kinesin-5 serves indispensable functions during mitosis, making an analysis of its function in interphase or in terminally differentiated cells difficult. Conditional mutations, such as temperature sensitive alleles, can overcome these limits of genetic analysis [3]. Gene knock down by RNAi approaches relays on protein turnover, leading to insensitivity of stable proteins. Pharmacological approaches with small molecules inhibitors or specific antibodies provide an alternative and have been applied for motor protein inhibition [4–6]. However, chemical approaches cannot be generalized, and need to be developed case by case.

Kinesin-5 belongs to kinesin family member 11 (KIF11), with the motor domain on N terminus, followed by a coiled-coil rod containing a central bipolar assembly (BASS) domain. Forming bipolar homo tetramers, Kinesin-5 can crosslink anti-parallel aligned microtubules. The motor activity enables filament sliding, e. g. during formation and elongation of the mitotic spindle [7]. In Drosophila syncytial embryos, Kinesin-5 is enriched at mitotic spindles and is essential for spindle formation and chromosome segregation. Injection of antibodies specific for Kinesin-5 into embryos leads to collapse of newly formed spindle and the formation of mono-asters of microtubules [5,6].

Making proteins susceptible to proteolytic cleavage represents a generally applicable strategy for generation of conditional alleles[8–10]. Here we apply this concept to motor proteins by inserting a proteolytic site between the head and stalk region (“neck”). We designated this strategy a “molecular guillotine” (Fig. 1A). We chose well-characterized Kinesin-5 in order to demonstrate the feasibility of this approach. As a protease, we employ TEV, which is highly specific. No match of TEV recognition motif within the Drosophila proteome has been identified, and flies expressing TEV are viable and fertile [10].

**Figure 1.**
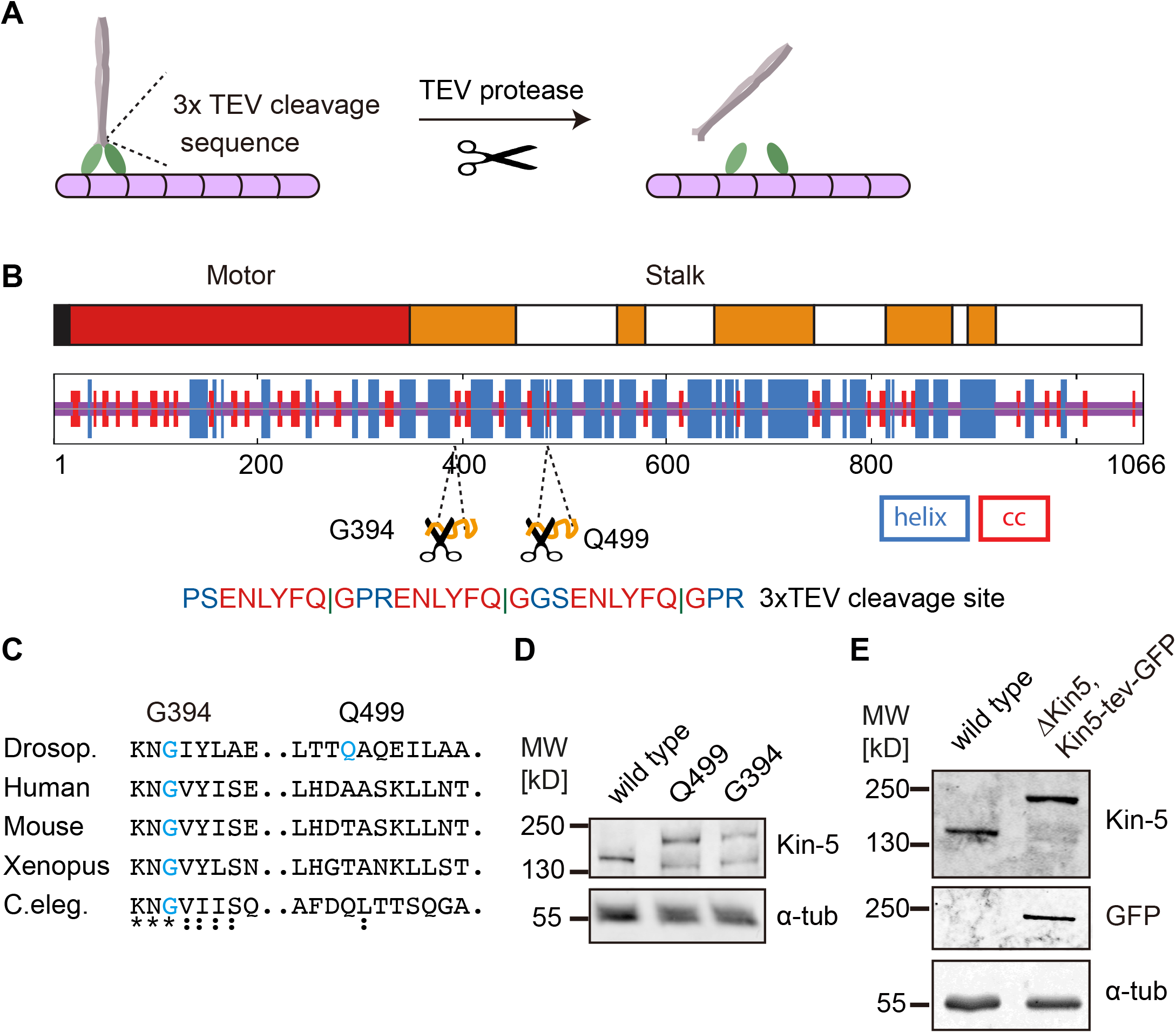
Design of a molecular guillotine for Kinesin-5. (**A**) Schematic illustration of motor protein molecular guillotine by inserting a protease substrate site next to the head domain of a motor. (**B**) TEV cleavage site in three copies (3x) is inserted in the coiled-coil region in stalk domain at position G394 or Q499. Domain structure of Kinesin-5 (Drosophila, Klp61F) and secondary structure prediction are indicated (**C**) Sequence alignment of the insertions sites at G394 and Q499 in wild type and Kinesin-5 mutant background. (**D, E**) Western blots with embryonic extracts (0-4h) from wild type and flies with the Kinesin-5-tev-GFP transgene, probed as indicated with antibodies for Kinesin-5, GFP, α-tubulin. Apparent molecular weight in kilo Dalton.

## Results

### Design of a “molecular guillotine”

We inserted three copies of the TEV recognition motif at one of two positions, G394 or Q499, into the stalk region. G394 and Q499 are located within conserved coiled-coil regions next to the head domain (Fig. 1B, C). In addition, we fused GFP to the C-terminus, which does not affect the function of Kinesin-5, as previously reported [11]. These constructs were expressed as transgenes in levels comparable to the endogenous allele with a ubiquitously active promoter, as assayed by western blot (Fig. 1D). Due to the C-terminal GFP moiety, the constructs showed a slower mobility in SDS-PAGE than wild type Kinesin-5. The TEV sites do not affect the functionality of Kinesin-5 as the construct with the insertion at G394 (Kin-5[G394tev]-GFP) complemented the lethality of a *Kinesin-5 (Klp61f)* mutation. For this, we recombined Kin-5[G394tev]-GFP with a *Kinesin-5* mutation. The resulting flies only expressed Kin-5[G394tev]-GFP, were viable and fertile and can be kept as a homozygous stock. In embryos from this line, Kinesin-5 was detected only at the molecular weight corresponding to transgenic Kin-5[G394tev]-GFP, which confirms the absence of endogenous Kinesin-5 (Fig. 1E).

### Kinesin-5 cleavage in vivo

We expressed TEV protease in stripes in embryos under the control of the *engrailed* promoter. Control embryos with no TEV expression showed uniform Kin-5[G394tev]-GFP expression. In contrast, the GFP signal was strongly depleted in stripes with TEV expression (Fig. 2A). Next we turned to syncytial embryos, which are characterized by their rapid and synchronous nuclear division cycles and the associated remodeling of the cytoskeleton. During mitosis, microtubules and their motors are important for formation and function of mitotic spindles and chromosome segregation, whereas they function in nuclear arrangement and stabilization of the nuclear array in interphase [12,13]. Kinesin-5 localizes to the mitotic spindle and is involved in chromosome segregation during mitosis [5,6,11]. We microinjected TEV protease into syncytial embryos and recorded GFP fluorescence. Following injection of TEV protease but not buffer, GFP fluorescence rapidly dropped (Fig. 2B). Correspondingly, the specific staining pattern, such as labelling of mitotic spindles or cytoplasmic asters was lost in TEV injected embryos (Fig. 2D). Quantification of total GFP fluorescence provided an estimate for an approximate half life of about 30 min (Fig. 2C). Kinesin-5 was specifically cleaved, since the electrophoretic mobility of Kin-5[G394tev]-GFP was higher in TEV than buffer injected embryos (Fig. 2E). Kin-5[G394tev]-GFP embryos were lysed about 30 min after injection and extracts analyzed by western blot against the C-terminus of Kinesin-5. The observed difference in electrophoretic mobility was consistent with proteolytic cleavage at the TEV sites at the neck and corresponding loss of the head domain. As we detected a single band, proteolytic cleavage was close to complete under our experimental conditions (Fig. 2E).

**Figure 2.**
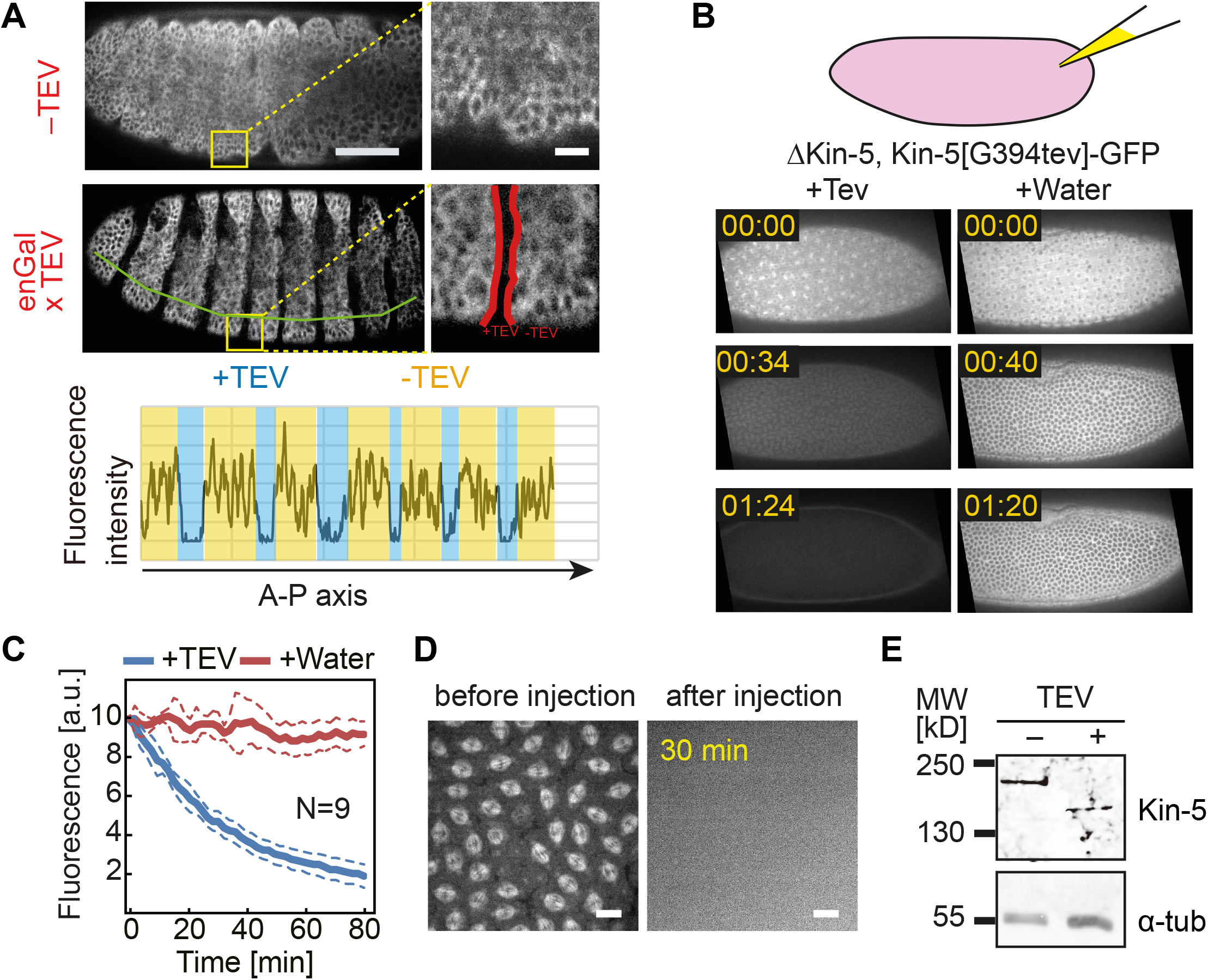
Kin-5[G394tev]-GFP is cleaved by TEV protease. (**A**) Image of living embryos expressing Kin-5[G394tev]-GFP with or without TEV protease expressed in striped pattern. Scale bar 50 μm. Region marked by squares in yellow are shown in high magnification. Scale bar 10 μm. Quantification of GFP signal along the anterior-posterior body axis (line in green). (**B-D**) TEV protease or buffer was injected into syncytial embryos mutant for *Kinesin-5* and expressing Kin-5[G394tev]-GFP. (**B**) Images from time lapse recording. Time in minute:second. (**C**) Quantification of GFP fluorescence. N, number or embryos. (**D**) Images of living embryos before and 30 min after injection. Scale bar 10 μm. (**E**) Western blot with extracts from embryos 30 min after injection with TEV or buffer probed with Kinesin-5 and α-Tubulin antibodies. Apparent molecular weight in kilo Dalton.

### Cleavage of Kinesin-5 leads the loss-of-function in mitosis

Next we analyzed the functional consequences of the Kinesin-5 cleavage. To track the nuclear cycles and behavior of chromosomes, we co-injected fluorescent labelled histone-1 and TEV protease into *Kinesin-5* null embryos expressing the Kin-5[G394tev]-GFP transgene. Following TEV injection, we observed a failure of chromosome separation and monoastral spindles (Fig. 3). These phenotypes were observed in individual spindles interspersed between normally appearing spindles. These phenotypes were consistent with the previously reported mitotic defects following Kinesin-5 antibody injection [5].

**Figure 3.**
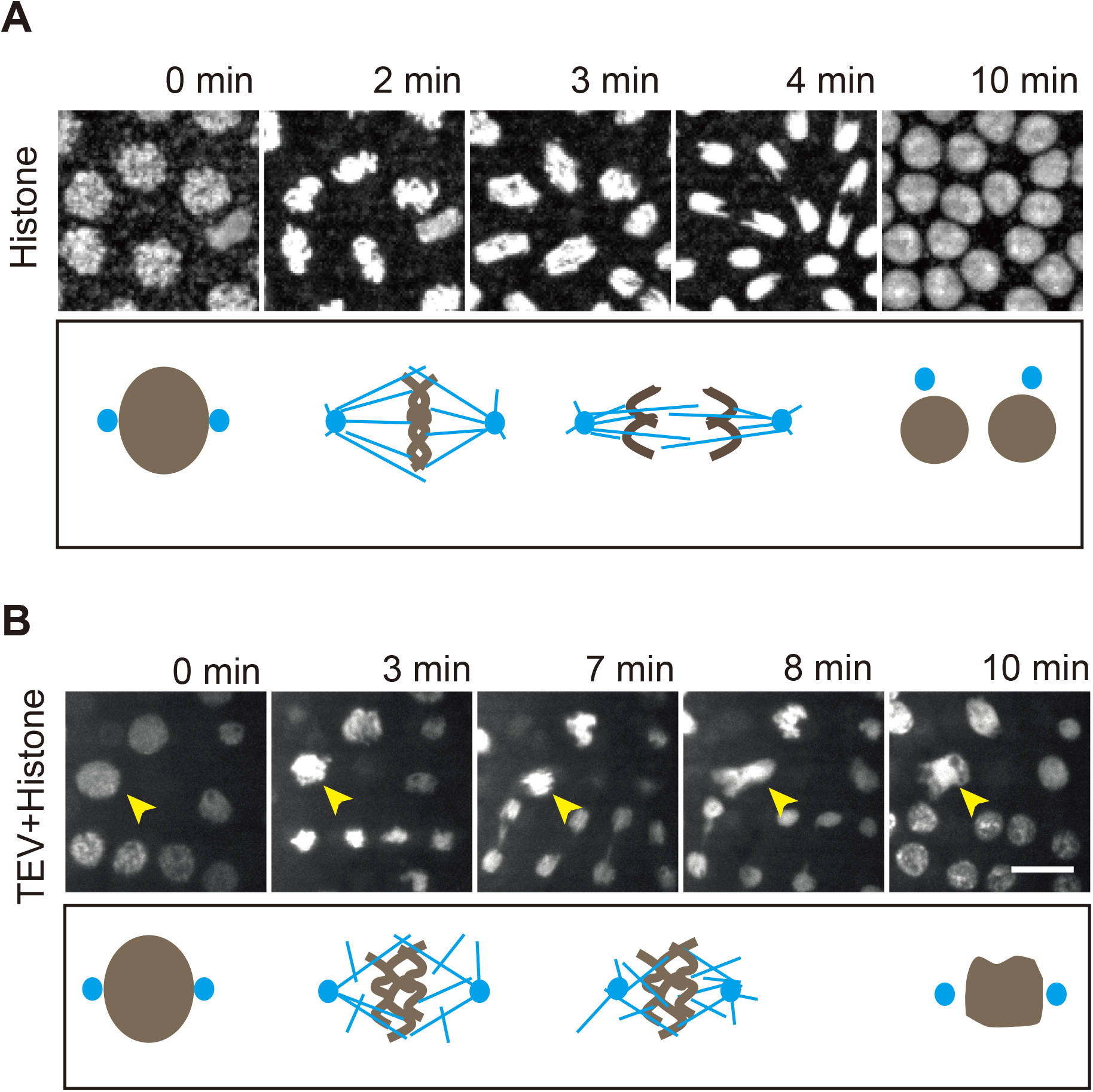
Phenotype of Kin-5[G394tev]-GFP cleavage by TEV protease. (**A, B**) Images from time lapse recording of embryos mutant for *Kinesin-5* expressing the Kin-5[G394tev]-GFP transgene and injected with fluorescent labeled Histone H1. (**B**) Coinjection of TEV protease. Arrow head in yellow points to defective mitotic figure. Schematic drawing of the mitotic figures. Scale bar: 10 μm.

### Interphase function of Kinesin-5

An interphase function of Kinesin-5 has not been investigated, yet. In interphases of syncytial embryos, Kinesin-5-GFP is strongly enriched at the centrosomes and associated asters. In addition, dynamic extended structures between adjacent asters were detected (Fig. 4C). These transient signals may represent microtubules coated with Kinesin-5 and possibly antiparallel aligned microtubules.

**Figure 4.**
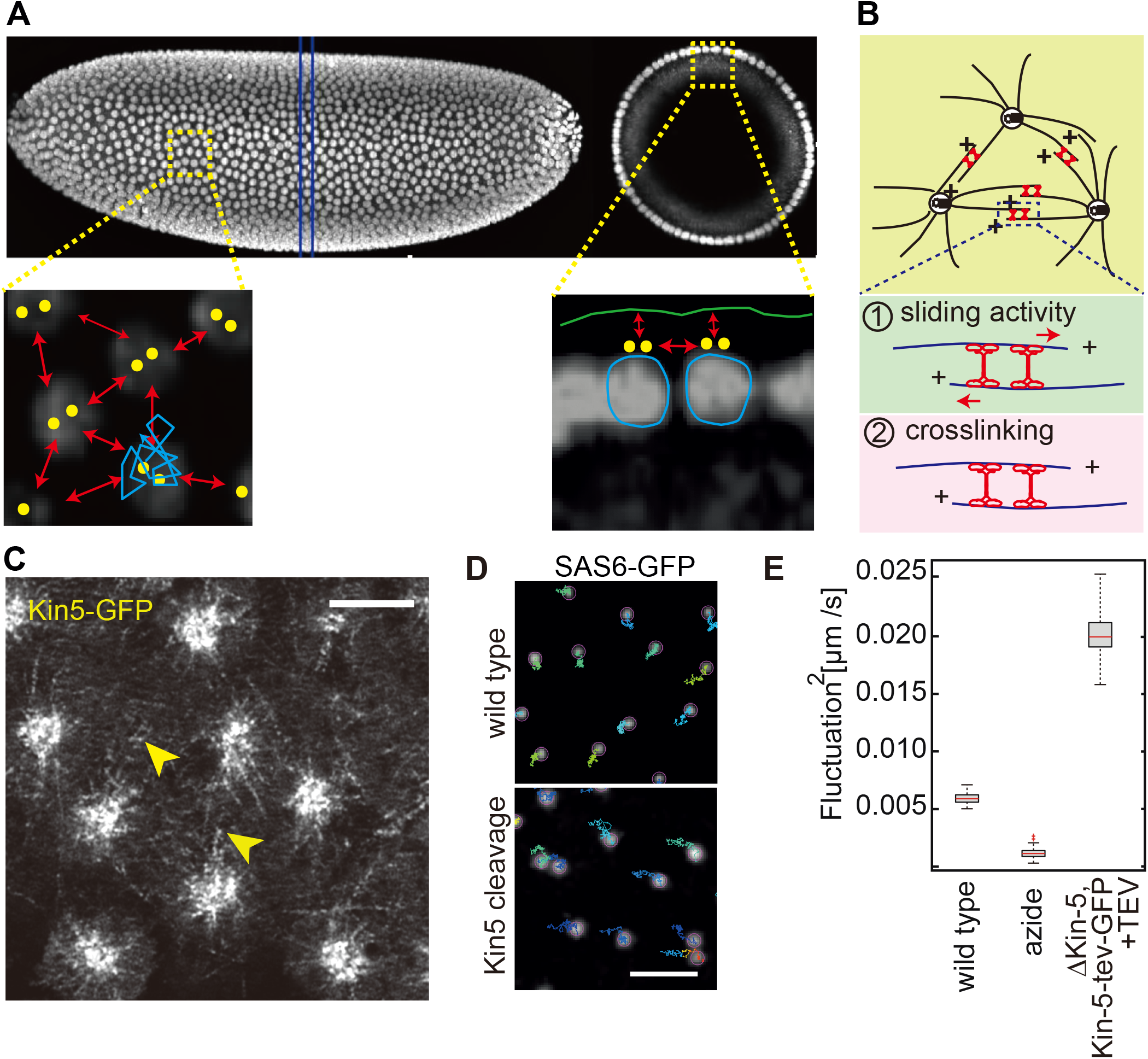
Interphase function of Kinesin-5. (**A**) Projected image of an embryo expressing Histone 2Av from selective plane illumination microscopy in side view and cross section (position indicated by lines in blue). Magnified section illustrate the interactions between the nuclei and between nuclei and cortex. Dots in yellow indicate centrosome pairs. (**B**) Illustration of microtubule asters with overlapping microtubules in anti-parallel orientation. Kinesin-5 may slide microtubules apart (Model 1) or crosslink adjacent asters (Model 2). (**C**) Image of living embryo expressing Kinesin-5-GFP (apical position) Scale bar 5 μm. (**D**) Images from living embryo mutant for *Kinesin-5* expressing Kin-5[G394tev]-GFP and SAS6-GFP and injection with TEV protease or buffer. Trajectories of centrosomes over 220 s on an image from time lapse recoding. Scale bar 5 μm. (**E**) Box plot displaying time-averaged fluctuation of centrosomes in embryos expressing SAS-6-GFP injected with buffer (wild type), sodium azide or TEV protease for cleavage of Kin-5[G394tev]-GFP.

As hypothesized previously [12,13], Kinesin-5 may be involved in nuclear positioning and formation of the nuclear array in syncytial Drosophila embryos. Kinesin-5 bound to anti-parallel aligned microtubules may push adjacent asters away from each other and thus generate a repulsive force, which may lead to uniform internuclear distances. In this model, Kinesin-5 would promote movements of centrosome and their associated asters. Alternatively, Kinesin-5 may crosslink microtubules from adjacent asters and stabilize the syncytial microtubule network. In this model Kinesin-5 would suppress movement of centrosomes and associated asters (Fig. 4B). To distinguish these two models, we recorded the dynamics of centrosomes in the scale of seconds [13]. From the recorded tracks, fluctuations of the centrosomes were calculated as previously reported [13]. These fluctuations have the dimension of a diffusion constant and do not contain slow (minute-scale) drift movement. We recorded centrosome dynamics in embryos with injected TEV protease and calculated the second-scale fluctuations (Fig. 4D). We find that the fluctuations are strongly increased to about 20x10^-3^ μm^2^/s as compared to about 6x10^-3^ μm^2^/s in control embryos injected with buffer. Passive fluctuations as detected in embryos depleted of ATP are in the range of 1.2x10^-3^ μm^2^/s [13] (Fig. 4E). Since cleavage of Kinesin-5 leads to an increased centrosome mobility, we conclude that functional Kinesin-5 stabilizes the dynamics of the microtubule array.

## Discussion

The function of Kinesin-5 for spindle formation and elongation during mitosis is well established [6]. Consistently, inhibition of Kinesin-5 by antibody injection induces defects in chromosome segregation in syncytial Drosophila embryos. Although expressed, a function of Kinesin-5 during interphase has been unknown, partly because such an interphase function was obscured by the mitotic defects in Kinesin-5 depleted embryos. The problem that one phenotype obscures other phenotypes is common to proteins with widespread functions, such as molecular motors. To circumvent this problem, we developed a method for conditionally inactivating Kinesin-5. In addition to Kinesin-5, this method is potentially suitable for other motor proteins, as well. With a “molecular guillotine”, we specifically inactivated Kinesin-5 by administration of TEV protease. In this way, we revealed an interphase function for the stabilization of the syncytial microtubule network. In syncytial embryos, the microtubule asters originating from centrosomes can directly interact with neighboring asters, since they are not physically separated by plasma membranes. These interactions lead to formation of an extended network covering the embryonic cortex. The phenotypic behavior of centrosomes and their associated nuclei reflect their intrinsic properties but also, as part of the network, the influences from the neighbors. Adjacent microtubule asters potentially interact via crosslinkers such as Feo/Ase1p, bundling proteins or motors with sliding activity, such as Kinesin-5. Here we tested the hypothesis that Kinesin-5 generates repulsive forces between adjacent astral microtubules in interphase. We expected that a loss of force generation would have led to a reduced mobility of the network and its nodes, the centrosomes. Using the fluctuations of centrosomes as an indicator of network dynamics, we rejected our hypothesis, because we measured an increased mobility of the centrosomes, when Kinesin-5 was inactivated. We interpret this data in that the in vivo function of Kinesin-5 as a crosslinker is more dominant than its function for sliding of anti-parallel aligned microtubules and thus pushing apart adjacent microtubule asters. The in vivo function of Kinesin-5 is similar to Kinesin-1, which is enriched at the cortex and F-actin and actin caps. Both may be involved in anchoring microtubule asters to the cortex and in this way counteract fluctuation movements of centrosomes. Having identified a suppressive function of Kinesin-5, the questions remains about the origin of the forces driving centrosome fluctuations. Fluctuations are due to an active component, since ATP depletion leads to loss of fluctuations. The (-)-end directed motor Kinesin-14 may serve as a force generator [6].

### “Molecular guillotine” can be used as a conditional mutant tool to study the fuction of motor protein

The “molecular guillotine” is potentially a versatile method for conditional inactivation of motor proteins. TEV protease has been used in inactivation of cohesin in yeast [14] and in fly [9], as well as Drosophila claudin [10]. However, this approach had not been used in motor proteins. The approach of a “molecular guillotine” as reported in this study can be applied widely to members of the motor protein families. Unlike using the small molecules inhibitor [4,15], TEV protease can be specifically expressed using UAS-GAL4 system in any genetically tractable cell type, and thus decapitate the selected motor protein in a tissue and developmental stage specific manner. Direct cleavage by TEV potentially leads to a faster inactivation kinetics than by the degron [16] or deGradFP systems [17], which rely on the ubiquitin-mediated protein degradation machinery. In addition, inactivation of motor protein by the “molecular guillotine” approach can be fine-tuned by titrating the TEV protease concentration, which help to identify additional functions of the motor proteins. In summary, the novel approach of a “molecular guillotine” enabled us to investigate a specific function of the motor protein Kinesin-5 in interphase. Potentially, the decapitation approach can be correspondingly applied to other kinesin motors as well as dyneins and myosins, as they have a related domain structure in common.

## Materials & Methods

### Genetics

Fly stocks (en-Gal4, Sas6-GFP, Klp61f ^07012^) [18,19] were obtained from the Bloomington Stock Center, if not otherwise noted. Transgenes of ubi-Kin5-tev-GFPQ499, ubi-Kin5-tev-GFPG394 and sqh-Kin5-GFP were generated by P element mediated random genome integration. We isolated multiple insertions on the third chromosome with varying expression. The ubi-Kin5-tev-GFPG394 line with strongest GFP fluorescence was recombined with an amorphic *Kinesin-5* mutation (Klp61f^07012^) and kept as a homozygous line. The transgenes spq-Kin5-GFP without TEV sites complemented the lethality of the *Kinesin-5* mutation *(Klp61f^07012^)* as well. TEV protease was expressed from a UASt-TEV transgene^9^ or injected as a purified recombinant protein.

### Cloning

A sequence coding for three recognitions sites of TEV protease (PS ENLYFQG PR ENLYFQG GS ENLYFQG PR) was inserted behind the codons of G394 or Q499 of the *Kinesin-5* cDNA. This sequence was cloned together with the ubiquitin promoter and eGFP into the multiple cloning site of a pUASt vector derivative lacking the UAS and hsp70TATA sites. *Kinesin5-GFP* was cloned by fusing eGFP coding sequence to the 3’ end of *Kinesin-5* coding sequence and transferred to transformation vector sGMCA [20]. Sequence information and details of the cloning procedure are available upon request.

### Western blotting, Kinesin antibody

The *Kinesin5* coding sequence corresponding to the C-terminal tail (aa 600-1066) was cloned by PCR with SK-Klp61f (Drosophila genomic resource center, Bloomington) as template into a protein expression vector with a N-terminal 9xHis tag. The His9-Kinesin-5-C600 protein with an apparent molecular weight of about 70 kd in SDS polyacrylamide gel electrophoresis (SDS-PAGE) was purified under denaturing conditions (Trenzyme, Konstanz) and used for immunization of rabbits (BioGenes, Berlin). Embryonic extracts were analyzed by SDS-PAGE and immunoblotting as previously described [21]. Briefly, proteins were blotted to nitrocellulose filters by wet transfer (100 mA per mini gel, overnight). The blots were blocked with 5% fat-free milk in PBS, incubated with primary antibodies in PBT (PBS with 0.1% Tween20, Kinesin-5, rabbit, 1:5000, α-Tubulin, mouse, 1:100000, B512, Sigma, GFP, rabbit, 1:5000, Torrey Pines Biolabs) and fluorescently labelled secondary antibodies (LiCOR, 1:20000, 0.05 μg/ml in PBT) for each two hours at room temperature. The developed blots were recorded with a LICOR system.

### Microinjection

Embryos were dechorionated and aligned on a coverslip, desiccated for 10 min, and covered with halocarbon oil (Voltalef 10S, Lehmann & Voss). We injected TEV protease at 10 μM purified from overexpressing E. coli (a gift from Dirk Görlich) or Histone1-Alexa-488 at 2 mg/ml (ThermoFisher).

### Microscopy

Images were recorded with a Zeiss microscope equipped with a spinning disc (25x/NA0.7 multi immersion, 40x/NA1.3oil). Centrosome movement was recorded in Sas6-GFP expressing embryos as previously described with a frame rate of 1 Hz [13]. Kin-5-GFP distribution in interphase was recorded with a confocal microscope (Zeiss LSM780 with airy scan unit, 63xNA1.4/oil). Images were processed with Fiji/ImageJ.

### Fluctuation analysis

The centrosomes tracking and measurement of fluctuation were carried out as previously described [13].

## Acknowledgements

We are grateful to D. Görlich, M. Gummalla, L. Henn, T. Kanesaki, D. Klopfenstein, R. Schuh for materials, preliminary experimental work or discussions. We acknowledge service support from the Developmental Studies Hybridoma Bank created by NICHD of the NIH/USA and maintained by the University of Iowa, the Bloomington Drosophila Stock Center (supported by NIH P40OD018537) and the Genomic Resource Center at Indiana University (supported by NIH 2P40OD010949-10A1). This work was in part supported by the Göttingen Centre for Molecular Biology (funds for equipment repair) and the Deutsche Forschungsgemeinschaft (DFG SFB937/TP10 and equipment grant INST1525/16-1 FUGG).

